# Multimodal Liquid Biopsy Improves Longitudinal Circulating Tumor DNA Detection in Children with Central Nervous System Tumors

**DOI:** 10.64898/2026.11.11.766035

**Authors:** Shea Gallus, Sharvari Mankame, Adriana Marshall, Ben Posorske, Alyssa Hernandez, Lisa Keller, Shari Kyman, Kazuhiro Sabet, Patricia Cornejo, Tania Contente-Cuomo, Jonathan Keats, Peter Chiarelli, Jason Hauptman, Michael E. Berens, Floris P. Barthel, Ross Mangum

## Abstract

**Background:** Circulating tumor DNA (ctDNA) is a promising biomarker for pediatric central nervous system (CNS) tumors, but limitations in sensitivity and universally accepted detection criteria remain. We evaluated multimodal ctDNA detection using a novel composite scoring framework integrating mutation, copy number, and fragmentomic analysis in longitudinal plasma and CSF samples from children with CNS tumors.

**Methods:** Tumor, blood and/or CSF samples were collected from pediatric CNS tumor patients. DNA was extracted from plasma, CSF, tumor, and whole blood. All samples underwent targeted and whole genome sequencing. Plasma and CSF samples were evaluated for ctDNA positivity—defined by a composite score ≥2—utilizing a tri-modal approach of whole genome sequencing for copy number alterations and fragmentomics and targeted sequencing.

**Results:** The cohort included 28 patients: medulloblastoma (39%), diffuse midline glioma (25%), high-grade glioma (18%), pilocytic astrocytoma (11%), and ependymoma (7%). A total of 140 plasma and 27 CSF samples were collected longitudinally. Combining all three modalities identified ctDNA in 70% of CSF and 17% of plasma samples, exceeding rates achieved by any single modality alone. Post-diagnosis plasma ctDNA positivity was significantly associated with worse progression-free and overall survival. CSF demonstrated significantly higher ctDNA signal intensity than plasma across all assay modalities.

**Conclusions:** A multimodal composite scoring approach improves longitudinal ctDNA detection in pediatric CNS tumors. Although CSF remains the most sensitive biofluid, plasma ctDNA reflects disease progression in select patients. These findings support integrated, multimodal liquid biopsy strategies for disease monitoring and highlight the need for prospective validation of standardized detection criteria.

**KEY POINTS:**

1. Multimodal profiling of longitudinal plasma and CSF samples improves circulating tumor DNA detection in pediatric CNS tumors.
2. Post-diagnosis plasma ctDNA positivity was associated with inferior progression-free survival despite the known challenges of plasma-based detection in pediatric CNS tumors.

**IMPORTANCE OF THE STUDY:** Pediatric brain tumors are among the most devastating childhood cancers, yet tools for non-invasive disease monitoring remain limited. Circulating tumor DNA (ctDNA) detected in blood and cerebrospinal fluid (CSF) offers a window into tumor biology without repeated surgery or biopsy. Here, we developed a multimodal scoring framework combining mutation analysis, copy number profiling, and DNA fragmentomics to standardize ctDNA detection across biofluids. Integrating all three methods improved detection beyond any single approach alone, and plasma ctDNA positivity was associated with significantly worse survival outcomes. These findings support the continued development of multimodal liquid biopsy as a clinical tool for disease monitoring in pediatric brain tumors.

## INTRODUCTION

The use of liquid biopsy for detection of circulating tumor DNA (ctDNA) in plasma and cerebrospinal fluid (CSF) from pediatric patients with central nervous system (CNS) tumors has gained considerable interest over the past decade. With either biofluid, ctDNA constitutes only the tumor-derived fraction of the total cell-free DNA (cfDNA), the remainder of which originates from non-malignant cells. Intriguing clinical applications include less invasive molecular characterization ^1–5^, real-time monitoring of treatment response ^6^, detection of minimal residual disease ^7^, and earlier identification of disease recurrence ^8^. Multiple proof-of-concept studies have demonstrated the feasibility of ctDNA detection in pediatric CNS tumors and have shown associations between ctDNA positivity and increased disease burden, treatment response, and disease progression ^8–13^. These encouraging findings, coupled with rapid advances in sequencing technologies and decreasing turnaround times, have accelerated efforts to incorporate liquid biopsy analyses into clinical trials and translational research initiatives.

Despite this progress, significant barriers continue to limit widespread clinical implementation. Pediatric CNS tumors frequently exhibit lower mutational burdens and reduced cellular turnover compared with many adult malignancies, creating challenges for sensitive detection ^1^. Furthermore, substantial heterogeneity exists across published studies regarding sample collection methods, biofluid selection, sequencing platforms, analytical pipelines, and thresholds used to define ctDNA positivity. As a result, side-by-side comparison of findings across studies and institutions remains difficult, confounding consensus regarding optimal testing methodologies. An additional obstacle specific to CNS tumors thought to limit the amount of ctDNA in the peripheral blood is the blood-brain barrier ^14,15^. While CSF is widely recognized as the most sensitive biofluid for ctDNA detection in CNS tumors ^1,12,16^, repeated longitudinal CSF collection remains invasive and is often impractical for routine disease monitoring, highlighting the continued need to improve the performance of plasma-based approaches.

Many sequencing methods have been utilized for ctDNA detection, including - but not limited to - digital droplet polymerase chain reaction (ddPCR) ^9,12^, targeted next-generation sequencing (NGS) ^3,17^, and whole-exome sequencing ^12^. With this data, different analysis methods have been used including copy number alteration (CNA) profiling ^8^, and DNA fragmentomic analysis ^18^. Each platform provides distinct biological information and possesses unique strengths and limitations with respect to sensitivity, specificity, cost, turnaround time, and applicability across tumor types. However, reliance on any single assay may fail to capture the full spectrum of tumor-derived cfDNA present within a liquid biopsy. Given these challenges, there is growing interest in combining ctDNA detection technologies to improve sensitivity and specificity, thereby leading to increased clinical utility ^19^. Combining NGS targeted mutation profiling with genome-wide CNA analysis and DNA fragmentomic characterization may provide a more comprehensive assessment of ctDNA than any individual platform alone.

In this study, a multimodal liquid biopsy analysis technique was applied to a cohort of pediatric patients with CNS tumors across longitudinal samples taken throughout their clinical course. Using paired tumor tissue, plasma, and CSF specimens, ctDNA detection was assessed through targeted sequencing, shallow whole-genome sequencing-based CNA profiling, and DNA fragmentomic analysis. The objective of this study was to evaluate the feasibility of longitudinal plasma and CSF sample collection in pediatric patients with CNS tumors and to assess the utility of comprehensive, multimodal profiling for improved longitudinal ctDNA detection.

## METHODS

### Patient Selection

Pediatric patients diagnosed with primary CNS tumors over a five-year period from 2021 to 2025 at Phoenix Children’s Hospital (PCH) were identified and prospectively enrolled. Patients with newly diagnosed or treatment-refractory/recurrent tumors across the spectrum of low- to high-grade histology and including localized or metastatic disease were eligible for study inclusion. Patients with non-CNS solid tumors or benign, non-tumor intracranial lesions were ineligible. Informed consent and assent discussions were completed prior to study enrollment, per Institutional Review Board policy.

### Sample Collection

Initial blood and CSF samples, when available, were collected at the time of study enrollment. Additional longitudinal specimens were obtained throughout treatment and during post-therapy surveillance. Sample collection was integrated into routine clinical care and occurred during scheduled laboratory evaluations, following venous access obtained for clinically indicated imaging studies, or during planned CSF procedures such as lumbar punctures or Ommaya reservoir taps. Plasma samples were collected approximately every three months for patients receiving active therapy and at clinically significant time points, including suspected disease progression or relapse. For patients who had completed therapy, plasma collection timing was paired with surveillance imaging evaluations. CSF samples were obtained only when clinically indicated procedures were being performed as part of standard patient care. No additional venipunctures, CSF procedures, or clinical encounters were performed solely for research purposes.

### Sample Processing

#### Blood

Venous blood samples at target volumes up to 20 mL, not exceeding 3 mL/kg of body weight per 24 hours, were collected in EDTA tubes and processed within one to three hours of collection. These were centrifuged at 1500 relative centrifugal force (rcf), for 15 minutes at 4 °C. Plasma was aliquoted into 1-2 mL samples. Two cell pellet aliquots were collected as 0.5-1 mL samples.

#### CSF

Cerebrospinal fluid samples at target volumes of 1-10 mL were collected by lumbar puncture, during surgical intervention, through an Ommaya reservoir tap, or via an external ventricular drain and placed into sterile naked tubes. Within 15 minutes, samples were centrifuged at 3000 rcf for 15 minutes at 4 °C. Supernatant was pipetted out and cryopreserved as 1-2 mL aliquots.

#### Storage

Plasma, cell pellet, and CSF samples were placed in a -80 °C freezer within 1 to 3 hours of collection and placed in the vapor phase liquid nitrogen (-196 ° C) for long term storage until time for future batched analysis.

### Sample Extraction and Prep

#### Cell free DNA (cfDNA)

cfDNA was extracted from 1 mL of CSF and 1 mL of plasma using the MagMAX Cell-Free DNA Isolation Kit (ThermoFisher, Waltman, MA USA) following the manufacturer’s procedure for lysis with Proteinase K. Vortex incubations during bead binding and elution was performed using a horizontal tube vortex adapter on a Vortex-Genie 2 Vortex. Eluted cfDNA was quantified using the Qubit 1x dsDNA High Sensitivity (ThermoFisher) and integrity was assessed using the High Sensitivity D1000 Screentape on the Agilent Tapestation 4200 (Agilent, Santa Clara, CA, USA). Libraries were prepared using the Watchmaker DNA Library Prep Kit (Watchmaker Genomics, Boulder, CO, USA) optimized for low input (1-3 ng) using Twist Bioscience’s UMI Adapter System (Twist Bioscience, South San Francisco, CA, USA), with UMI adapters diluted to 3 µM and amplified using UDI primers for 13 cycles. Final libraries were quantified using Qubit 1X dsDNA High Sensitivity Kit and the average library size acquired using High Sensitivity D1000 Screentape. Libraries were then sequenced to a target of 3-5X whole genome coverage on the Illumina Novaseq X Plus platform using 2x150 paired end sequencing.

#### Genomic DNA (gDNA)

For each patient sample, formalin-fixed paraffin-embedded (FFPE) tumor tissue and whole blood (utilized as the “normal” non-tumor reference in subsequent analyses) were collected. DNA from the whole blood was isolated using Monarch HMW DNA Extraction kit for Cells & Blood (New England Biolabs, Ipswich, MA, USA). DNA from FFPE tissue was isolated using the QIAamp DNA FFPE Tissue Kit (QIAGEN, Hilden, Germany). DNA was quantified with Qubit 1X dsDNA Broad Range kit (ThermoFisher) and integrity was assessed using Genomic DNA ScreenTape (Agilent). Libraries were prepared using Watchmaker DNA Library Preparation with Fragmentation, following manufacturer’s recommendations with Twist Bioscience UMI Adapter System. Given the already fragmented nature of FFPE-derived DNA, library preparation protocols were adjusted by reducing fragmentation time to prevent over-shearing and increasing PCR cycles to compensate for the typically low DNA yields from FFPE samples.

#### Target Enrichment and Sequencing

Target enrichment was carried out using the Twist Target Enrichment Fast Hybridization Protocol (Twist Biosciences). Pools of up to eight libraries were enriched with both a mitochondrial panel and a custom glioma panel containing frequently mutated glioblastoma genes (Twist Biosciences). Libraries were combined at equal concentrations to generate a 1.5 µg pool, which was dried using a vacuum centrifuge. The pooled libraries were hybridized with capture panels and incubated at 60 °C for 4 hours. Enriched fragments were captured with streptavidin-coated beads, washed with supplied buffers, and eluted in 45 µl of molecular-grade water. Final libraries were quantified using Qubit and Tapestation. Genomic DNA libraries and cfDNA libraries were sequenced on the Illumina Novaseq X Plus platform using 2x150 paired end sequencing.

### Data Analysis

#### Bioinformatics Analysis

Raw sequenced reads were adapter-trimmed and aligned to GRCh38 (hg38). PCR duplicates were collapsed by UMI-aware consensus calling using fgbio’s standardized pipelines. Copy number and tumor fraction were estimated from low-pass whole-genome data using ichorCNA with 1 Mb bins. Arm-level aneuploidy was scored from the ichorCNA copy-ratio segments. Somatic SNVs were called from the targeted sequencing data using GATK Mutect2, then annotated and filtered to remove variants present in germline references. Tumor specific variants were identified by filtering out variants present in the patient’s matched germline.

#### Survival Analysis

Survival analyses were performed to evaluate the association between longitudinal ctDNA detection and patient outcomes. To focus on the prognostic value of longitudinal monitoring, samples collected on the day of diagnosis were excluded from ctDNA classification. Patients with at least one ctDNA-positive post-diagnosis plasma sample were classified as ctDNA-positive, whereas patients with no detectable ctDNA-positive post-diagnosis plasma samples were classified as ctDNA-negative. Patients without a post-diagnosis plasma sample were excluded from this analysis (n = 2), leaving 26 patients (7 ctDNA-positive, 19 ctDNA-negative). Results were similar when day-0 samples were included (**Table 1**). Overall survival was defined as the interval from diagnosis to death or last clinical follow-up. Progression-free survival was defined as the interval from diagnosis to radiographic evidence of tumor progression or recurrence. Survival distributions were estimated using the Kaplan–Meier method and compared between ctDNA-positive and ctDNA-negative groups using the log-rank test.

**Table 1.**
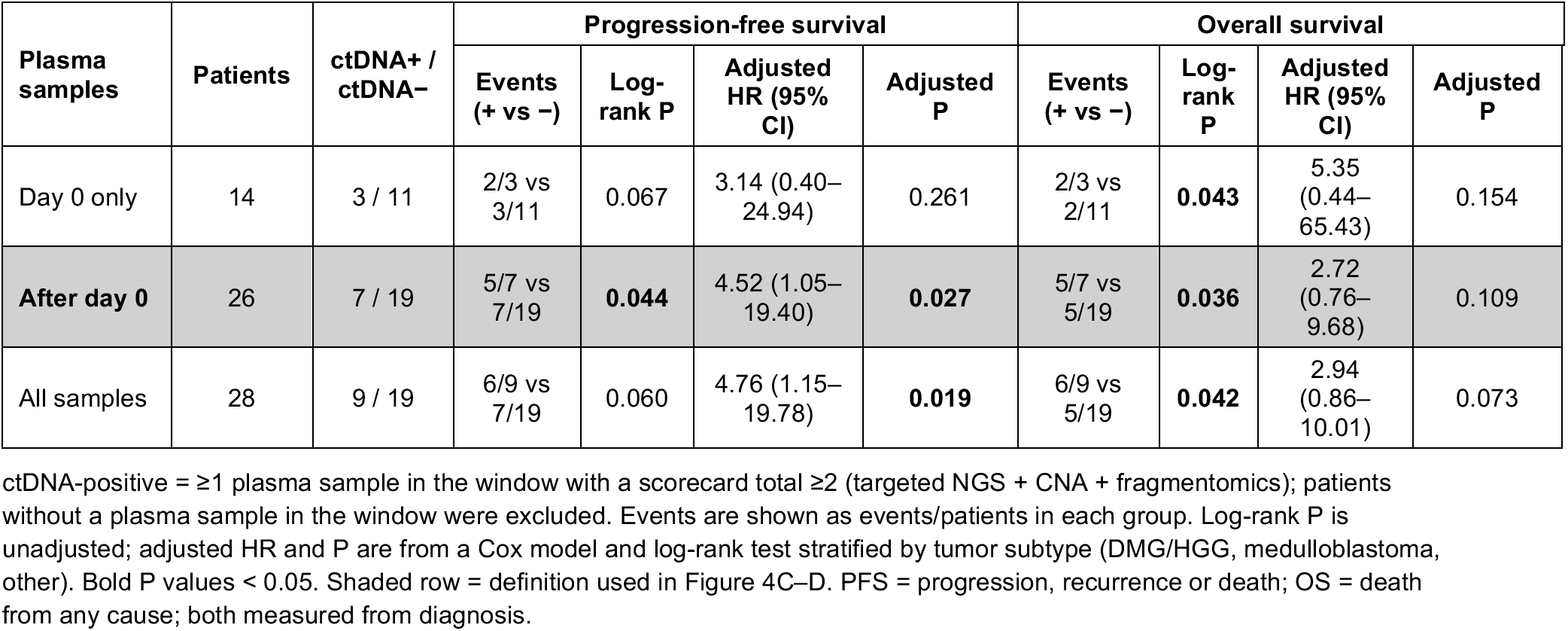
PFS and OS adjusted and unadjusted p values.

| Plasma samples | Patients | ctDNA+ / ctDNA− | Progression-free survival |  |  |  | Overall survival |  |  |  |
|---|---|---|---|---|---|---|---|---|---|---|
|  |  |  | Events (+ vs −) | Log-rank P | Adjusted HR (95% CI) | Adjusted P | Events (+ vs −) | Log-rank P | Adjusted HR (95% CI) | Adjusted P |
| Day 0 only | 14 | 3 / 11 | 2/3 vs 3/11 | 0.067 | 3.14 (0.40–24.94) | 0.261 | 2/3 vs 2/11 | **0.043** | 5.35 (0.44–65.43) | 0.154 |
| **After day 0** | 26 | 7 / 19 | 5/7 vs 7/19 | **0.044** | 4.52 (1.05–19.40) | **0.027** | 5/7 vs 5/19 | **0.036** | 2.72 (0.76–9.68) | 0.109 |
| All samples | 28 | 9 / 19 | 6/9 vs 7/19 | 0.060 | 4.76 (1.15–19.78) | **0.019** | 6/9 vs 5/19 | **0.042** | 2.94 (0.86–10.01) | 0.073 |

ctDNA-positive = ≥1 plasma sample in the window with a scorecard total ≥2 (targeted NGS + CNA + fragmentomics); patients without a plasma sample in the window were excluded. Events are shown as events/patients in each group. Log-rank P is unadjusted; adjusted HR and P are from a Cox model and log-rank test stratified by tumor subtype (DMG/HGG, medulloblastoma, other). Bold P values < 0.05. Shaded row = definition used in Figure 4C–D. PFS = progression, recurrence or death; OS = death from any cause; both measured from diagnosis.

#### Statistical Analysis

To compare ctDNA detection rates between plasma and CSF, patients with samples from both biofluid types were included in paired analyses (n=15). Differences in the proportion of ctDNA-positive patients between CSF and plasma were assessed using the McNemar test across four timepoint windows: at diagnosis (day 0), after diagnosis but before treatment initiation, at any post-diagnostic timepoint, and across all samples from day 0 onward. For quantitative comparison of signal intensity between biofluids, per-patient maximum values were used for each assay metric. Paired within-patient differences between CSF and plasma were assessed using the Wilcoxon signed-rank test. All statistical tests were two-sided, and P < 0.05 was considered statistically significant.

#### Scoring Analysis

To classify ctDNA detection, we developed a multimodal scoring system in which individual scores from all three detection methods (criteria summarized below) are summed into a single composite score. This combined metric determines the final ctDNA classification: a composite score of 0 indicates ctDNA-negative samples, a score of 1 is considered indeterminate due to insufficient evidence, and a score of ≥2 establishes positive ctDNA detection.

#### Variant Allele (Targeted NGS) Scoring Metrics

A tumor-informed approach was utilized to detect and quantify ctDNA from targeted sequencing data. Somatic variants were identified by comparing variant calls from matched tumor tissue and germline samples, retaining only variants present in the tumor but absent from the germline. A minimum read depth of 10-30x was required across all variant sites to ensure reliable variant detection and minimize false positives arising from sequencing noise. cfDNA samples were genotyped against the tumor specific variants, and the variant allele frequency (VAF) was calculated by the number of supporting reads (ALT) of that variant divided by read depth (DP). The scoring threshold for VAF was assigned after quantifying the baseline of noise through whole blood pellet (WBP) samples (**Supplemental Figure 2A**). A score of 0 was given if no mutations were detected in longitudinal samples. A score of 1 was assigned when a VAF of >0% to <1.0% was detected in variants not expected to alter protein function (e.g., intronic, RNA, synonymous, or other non-coding variants). A score of 2 was assigned when the VAF was ≥1.0% for variants not expected to alter protein function or when a pathogenic variant was detected at any VAF.

#### Copy Number Scoring Metrics

Copy number-based tumor fraction was estimated using ichorCNA ^20^, which models genome-wide read depth from WGS data to infer large-scale copy number alterations and calculate a tumor fraction estimate. To minimize false positive signal arising from germline copy number variants, each patient’s matched WBP was used as a patient-specific panel of normals, ensuring that only somatic copy number alterations contributed to the tumor fraction estimate. A score of 0 was given if tumor fraction was between 0% and 3%. A score of 1 was given if tumor fraction was ≥3 but less than 5 and there was >3% difference from the WBP fraction. A score of 2 was given if tumor fraction was >5% or if there was ≥3% difference from the WBP fraction. These values were chosen based on the threshold we observed in the WBP samples (**Supplemental Figure 2B**).

#### Fragmentation Scoring Metrics

Insert sizes for each fragment in the cfDNA and gDNA samples were calculated using Picard CollectInsertSizes. Short fragments were defined as 100–150 base pairs and long fragments as 151–220 base pairs ^21^. Positivity was defined as a short-to-long fragment ratio exceeding both the matched WBP ratio plus two standard deviations. A score of 0 was given if there was <2 standard deviations difference from the WBP. A score of 1 was given if there was >2 standard deviation difference from the WBP (**Supplemental Figure 2C**).

## RESULTS

### Characterizing a clinically diverse cohort of 28 pediatric CNS tumor patients

Fifty-seven patients with pediatric CNS tumors were enrolled on our ctDNA study from January 2021 through December 2025 **(Figure 1)**. From these patients, a pilot cohort of 28 patients (140 plasma samples and 27 CSF samples) was identified for further analysis. An intentional effort was made to select patients of varied histologic diagnoses with a combination of localized and metastatic disease. Most blood samples were collected either in the oncology clinic (n=65, 39%) during routine clinic visits with planned lab draws, or in the radiology department after the placement of a peripheral IV or central line access for planned MRI imaging (n=64, 38%). Other opportunities for blood sample collection included the operating room (n=28, 17%) and/or during inpatient admissions (n=10, 6%). CSF collections occurred during surgical resection (n=11, 41%) and lumbar punctures (n=10, 37%), or at the time of external ventricular drain (n=4, 15%) or Ommaya catheter (n=2, 7%) access (**Table 2**). Samples were taken at a variety of timepoints throughout the patient’s clinical course including at time of diagnosis (n=48, 29%), during active therapy (n=54, 32%), off-therapy surveillance (n=42, 25%), and/or at time of progression or relapse (n=23, 14%). Seventy of 140 (50%) plasma samples were taken in the setting of radiographic evidence of active disease at the time of sample collection compared to 20/27 (74%) of CSF samples. Five patients had more than one CSF longitudinal sample collected, ranging from 2-3 samples. The clinical scenarios for repeat CSF collections included two at initial staging, three at time of disease progression, and one Ommaya tap for intraventricular chemotherapy.

**Figure 1.**
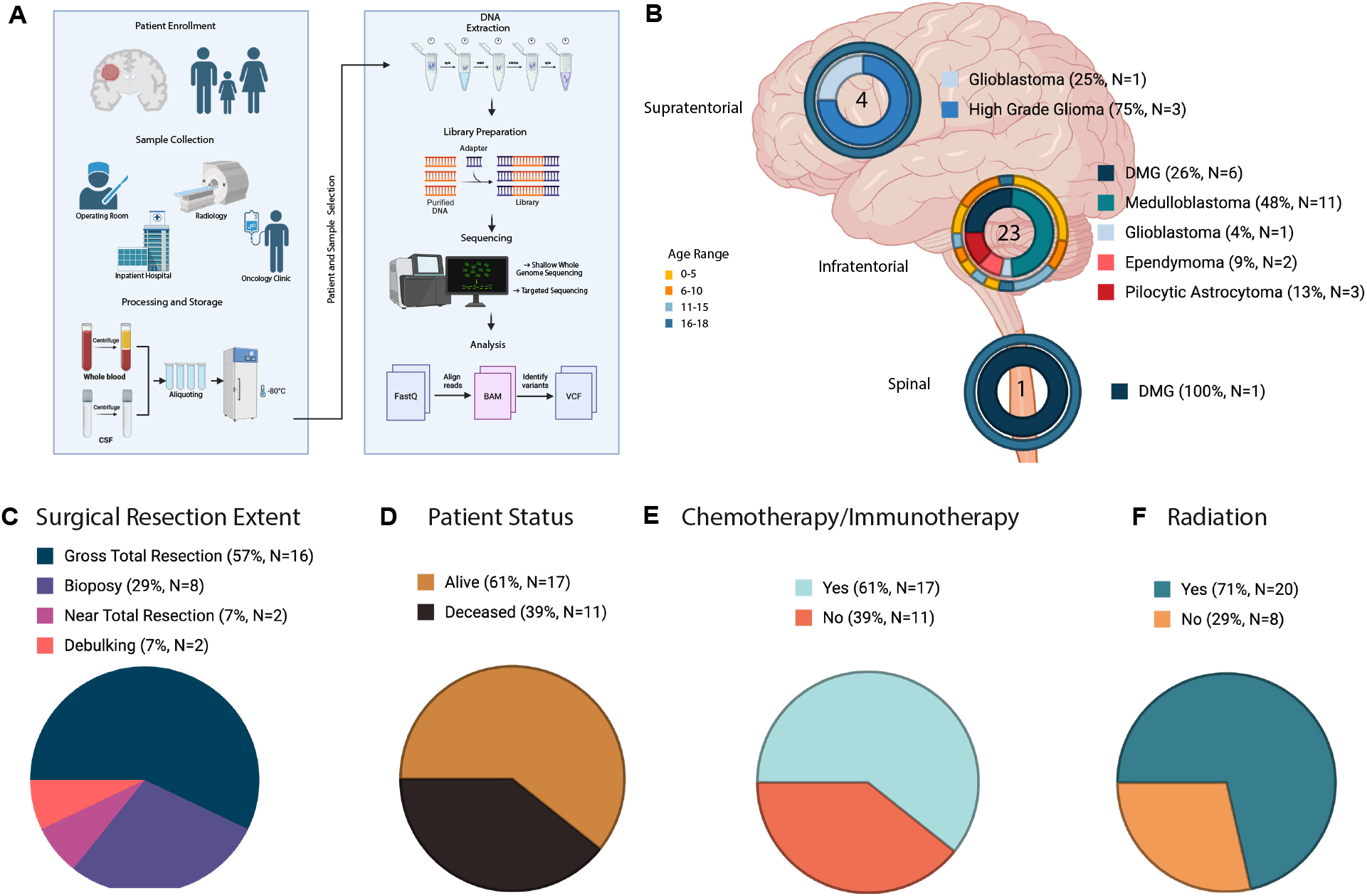
Overview of ctDNA Analysis Workflow and Patient Characteristics: **A.** Schematic of the workflow from patient enrollment to data analysis for the 28 patients included in this dataset. **B**. Tumor location categorized as supratentorial, infratentorial, or spinal. Number in the center of the circle is the total number of patients with tumors in that specific location. Inner circle indicates tumor diagnosis. Outer circle indicates age of patients. **C-F.** Pie charts of surgical resection extent, patient status, chemotherapy/immunotherapy, and radiation respectively. Created with BioRender.com.

**Table 2.** cfDNA Sample Collection Details.

|  | **CSF** | **Plasma** | **Total** |
|---|---|---|---|
| **Total number of samples** | 27 | 140 | 167 |
| **Average samples per patient** | 0.9 | 5 | 5.9 |
| **Average Day of Collection from Diagnosis (Days)** | 79.3 | 301 | 265 |
| **Range (Days)** | 0-749 | 0-993 |  |
| **Collection location** |  |  |  |
| **Clinic** | 10 | 55 | 65 |
| **Radiology** | 0 | 64 | 64 |
| **Operating Room** | 14 | 14 | 28 |
| **Hospital** | 3 | 7 | 10 |
| **Collection Timepoint** |  |  |  |
| **Presentation** | 21 | 69 | 48 |
| **Monitoring on Therapy** | 1 | 53 | 54 |
| **Monitoring off Therapy** | 1 | 41 | 42 |
| **Relapse/Progression** | 4 | 19 | 23 |
| **Method of CSF Collection** |  |  |  |
| **Lumbar Puncture** | 10 |  |  |
| **External Ventricular Drain** | 4 |  |  |
| **Surgical Resection** | 11 |  |  |
| **Ommaya Catheter** | 2 |  |  |

There was a slight female predominance (n=16, 57%) represented in the cohort, and patients were aged 1-18 years old (average 9.3 years) at the time of initial diagnosis. Tumors represented in the cohort included medulloblastoma (n=11, 39%), diffuse midline glioma (DMG) (n=7, 25%), high-grade glioma (n=5, 18%), pilocytic astrocytoma (n=3, 11%), and ependymoma (n=2, 7%) (**Figure 1B**).

Most tumors were infratentorial (n = 23, 82%), including all medulloblastomas, pilocytic astrocytomas, ependymomas, and six of seven DMGs. Four tumors (14%) were supratentorial, comprising three high-grade gliomas and one glioblastoma, and one DMG (4%) was in the spinal compartment (**Figure 1B**). Gross total resection was achieved in the majority of patients (n = 16, 57%), while biopsy was performed in 8 (29%), near total resection in 2 (7%), and debulking in 2 (7%) (Figure 1C). At last clinical follow-up, 17 patients (61%) were alive and 11 (39%) had died (**Figure 1D**). Over the course of their treatment, 17 patients (61%) received chemotherapy and/or immunotherapy (**Figure 1E**), and 20 (71%) received radiation therapy (**Figure 1F**). Of the 167 samples collected, 159 (95.2%) underwent successful cfDNA extraction, including 139 plasma samples and 20 CSF samples, and were used for subsequent analysis.

### Tumor genomic profiling reveals diagnosis-specific mutational landscapes and chromosomal instability in high-grade tumors

Targeted sequencing of 28 tumors identified 58 somatic alterations across 32 genes, with at least one alteration in 24 of 28 (86%) tumors (**Figure 2**). Most alterations were missense (n = 32), frameshift (n = 7) or nonsense (n = 1) variants; the remainder were non-coding (intronic, untranslated region, or flanking; n = 17). The most frequently mutated genes were *H3-3A* (n=7), *TP53* (n=7), *ATRX* (n=3), *PIK3CA* (n=4), *RB1* (n=2), and *SMARCA4* (n=3), with an average of 2.7 mutations per tumor and a mean variant allele frequency of 35.4% (SD 27.7%). Two patients with medulloblastoma did not have any tumor specific mutations identified on the targeted gene panel.

**Figure 2.**
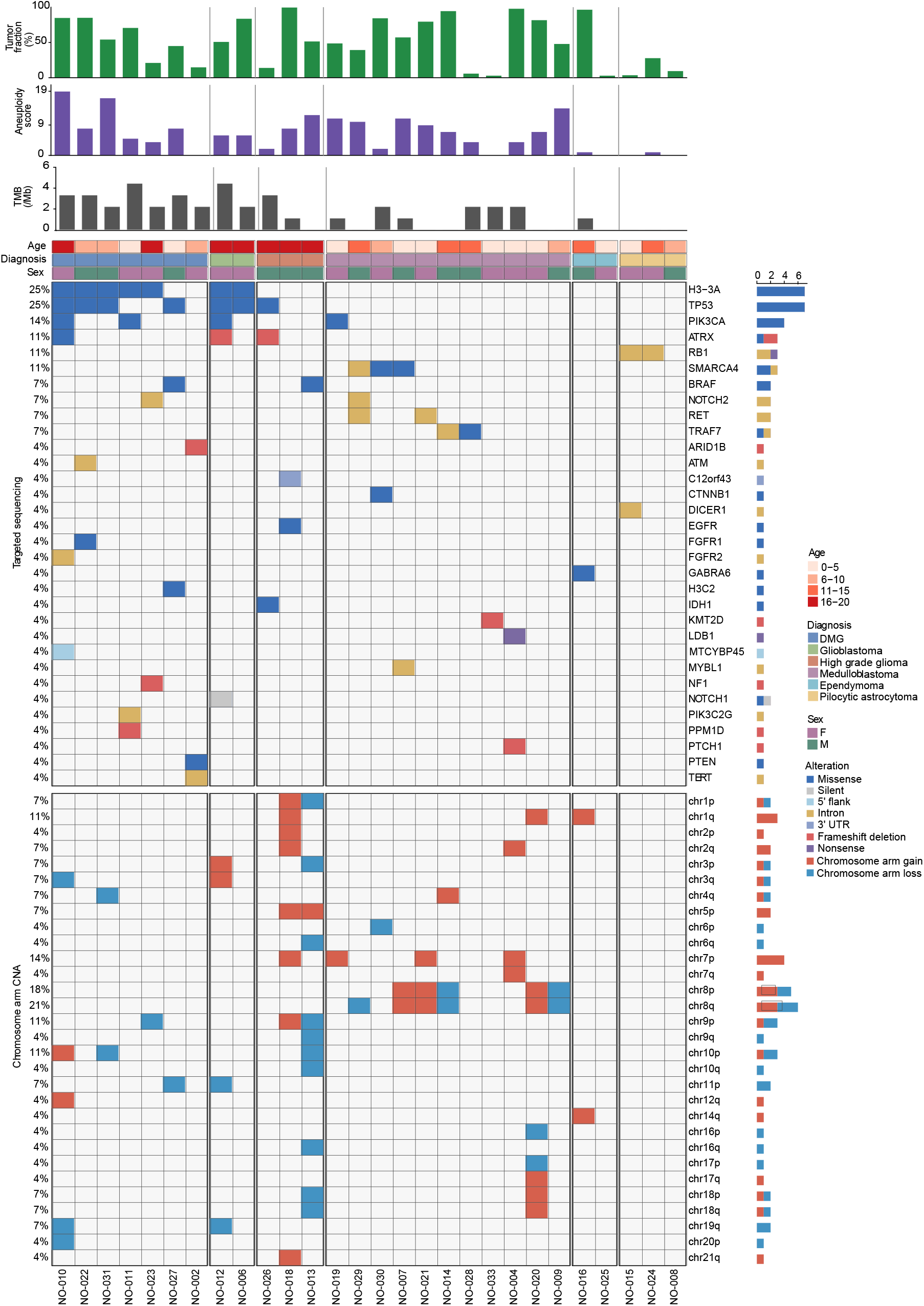
Oncoprint of Genomic Alterations in Study Cohort: Top-Bar plots show tumor fraction, aneuploidy score and tumor mutational burden (TMB) for each patient tumor. Columns represent individual patients and rows represent different genes with recurrent alterations. Samples are grouped by diagnosis. Mutation alterations are shown by color, illustrating the distribution and co-occurrence of genetic alterations across tumor types. Bottom-Chromosomal arm-level copy-number alteration heatmap. Red indicates copy-number gain, blue indicates copy-number loss, and white indicates copy-number neutrality.

The mutational landscape differed markedly by diagnosis. DMGs were the most heavily mutated group, with 25 alterations across 7 tumors; six of seven harbored an H3 K27M mutation (*H3-3A*, n = 5; H3C2, n = 1), frequently co-occurring with *TP53* (4/7) and *PIK3CA* (2/7). Both glioblastoma patients carried *H3-3A* K27M and *TP53* mutations. The three remaining high-grade gliomas showed distinct profiles without histone mutations, including *IDH1* R132G with *TP53* and *ATRX* frameshift deletion, *BRAF* V600E, and *EGFR* missense. Among medulloblastomas, 9 of 11 tumors harbored alterations enriched for known subgroup-associated genes, including *SMARCA4* (3/11), *CTNNB1*, and *PTCH1*. Low-grade and ependymal tumors showed few alterations, limited largely to non-coding *RB1* and *DICER1* variants.

Chromosomal instability was concentrated in high-grade tumors. Arm-level copy number changes were detected in 16 of 23 (70%) high-grade tumors compared with 1 of 5 (20%) low-grade or ependymal tumors (p = 0.06, Fisher’s exact test), and aneuploidy scores were significantly higher in high-grade tumors (median 7 vs. 0; p = 0.003, Mann-Whitney U test). High-grade gliomas carried the greatest per-tumor burden, with up to 11 arm-level events in a single tumor. Medulloblastomas exhibited recurrent, subtype-characteristic copy number patterns, including chromosome 7p gain and an isochromosome 17q-like pattern. Pilocytic astrocytomas had no arm-level events. Consistent with these findings, median tumor fraction was 54% in high-grade tumors compared with 10% in low-grade tumors (p = 0.10).

### Integration of Mutation, Copy Number, and Fragmentomic Data into a Unified Scoring System Improves ctDNA Detection

To systematically integrate plasma or CSF-derived ctDNA signal from all three modalities, each longitudinal sample was assigned a composite score based on contributions from targeted sequencing, CNA analysis, and fragmentomics (**Figure 3A, Supplemental Figure 1-2**). Samples scoring ≥ 2 were classified as ctDNA-positive. Across all 159 longitudinal samples, 38 (24%) were ctDNA-positive by this criterion (**Figure 3B)**.

**Figure 3.**
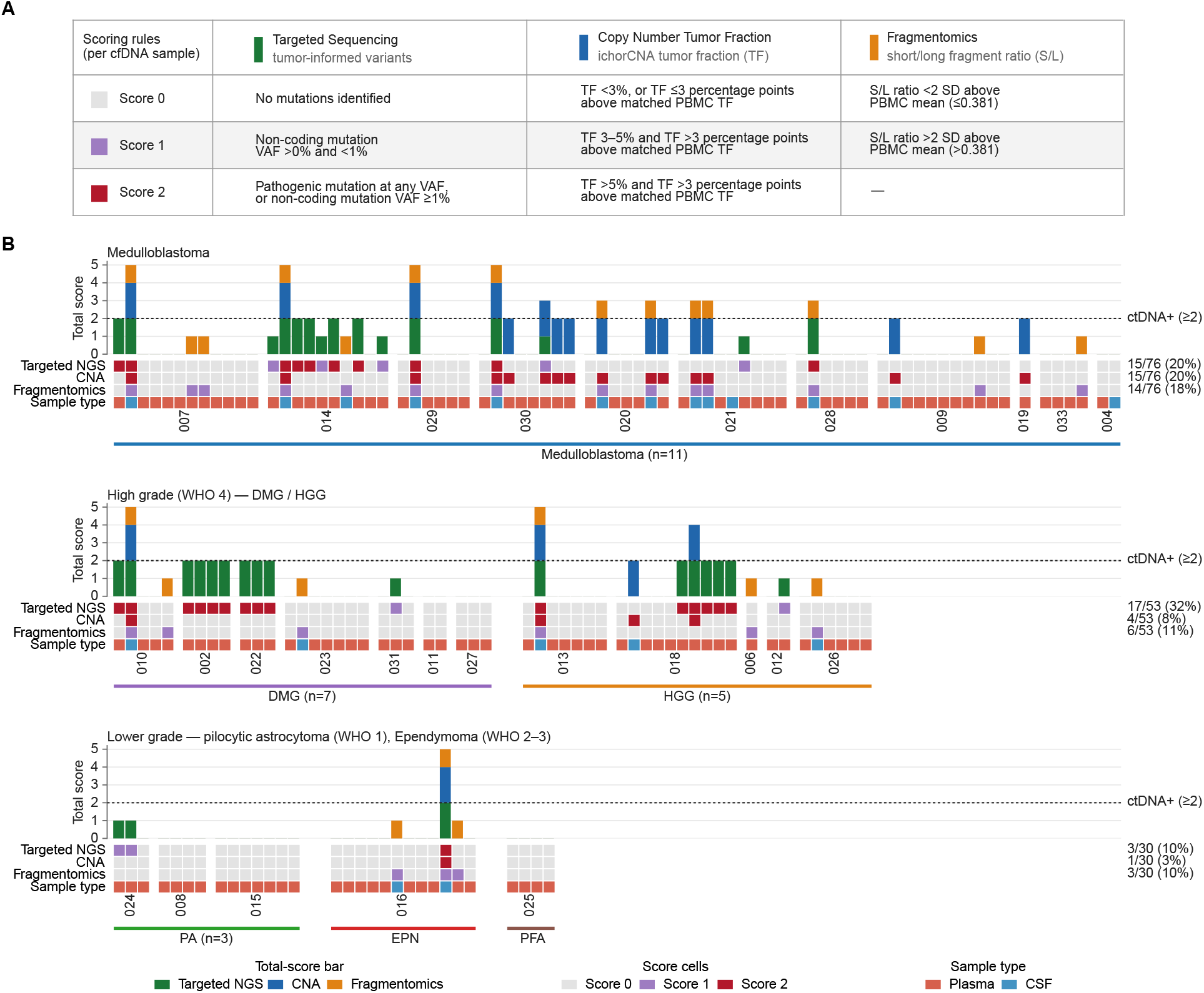
Oncoprint of Scoring Data Across Dataset. **A.** Table showing scoring rules separated by metric. Each sample receives a total score of 0-5 and scores of ≥ 2 are ctDNA positive. **B.** Each column represents a cfDNA sample (n=159; 139 plasma, 20 CSF) from 28 patients. Samples are ordered by collection day within patient, and patients are grouped by diagnosis and WHO grade: medulloblastoma; high grade (WHO 4) DMG and HGG; and lower grade (pilocytic astrocytoma, WHO 1; ependymoma and PFA ependymoma, WHO 2–3). Cells are colored in terms of score with grey (0), light red (1) or dark red (2). Bars show each sample’s total score (0–5), colored by assay contribution. Samples with a total ≥ 2 (dashed line) were called ctDNA-positive (55/167, 33%). Numbers at right give the samples scoring > 0 for each assay within.

Among medulloblastoma patients (76 samples), targeted sequencing (15/76, 20%) and CNA (15/76, 20%) were the most frequently positive assays, followed by fragmentomics (14/76, 18%). In DMG and HGG patients (53 samples), targeted sequencing yielded the highest positivity rate (17/53, 32%), with CNA (4/53, 8%) and fragmentomics (6/53, 11%) providing complementary signal. Low-grade and ependymal tumors showed markedly lower positivity across all three methods (NGS 3/30, 10%; CNA 1/35, 3%; fragmentomics 3/30, 10%), consistent with their lower tumor burden and reduced genomic instability.

The thresholds applied within each scoring tier were empirically derived and validated using matched germline samples as a negative reference (**Supplemental Figure 2**). For variant allele scoring, no germline samples harbored pathogenic or non-coding variants at VAF ≥1% (0% for both classes, n=36 and n=38 respectively), confirming that the 1% threshold effectively excludes germline background (**Supplemental Figure 2A**). For CNA scoring, ichorCNA tumor fraction estimates in germline samples ranged from 0–4.1% (median 3.1%), with no samples exceeding 5%, supporting the 5% tumor fraction cutoff as a clean boundary above germline noise (**Supplemental Figure 2B**). For fragmentomic scoring, the short-to-long fragment ratio cutoff of 0.381 was defined as the germline mean plus two standard deviations (mean 0.223) and no germline samples exceeded this threshold. Using this cutoff, 43% of tumor samples and 80% of CSF samples fell above the threshold, compared to only 5% of plasma samples (**Supplemental Figure 2C**).

### ctDNA positivity is significantly associated with worse progression free survival and overall survival

Across all three metrics, 15 out of 28 patients (54%) had at least one tumor positive CSF or plasma sample (**Figure 4A**). ctDNA was detected significantly more frequently in CSF samples (14/20, 70%) than in plasma samples (24/139, 17.3%; Fisher’s exact test, *P* < 0.0001).

**Figure 4.**
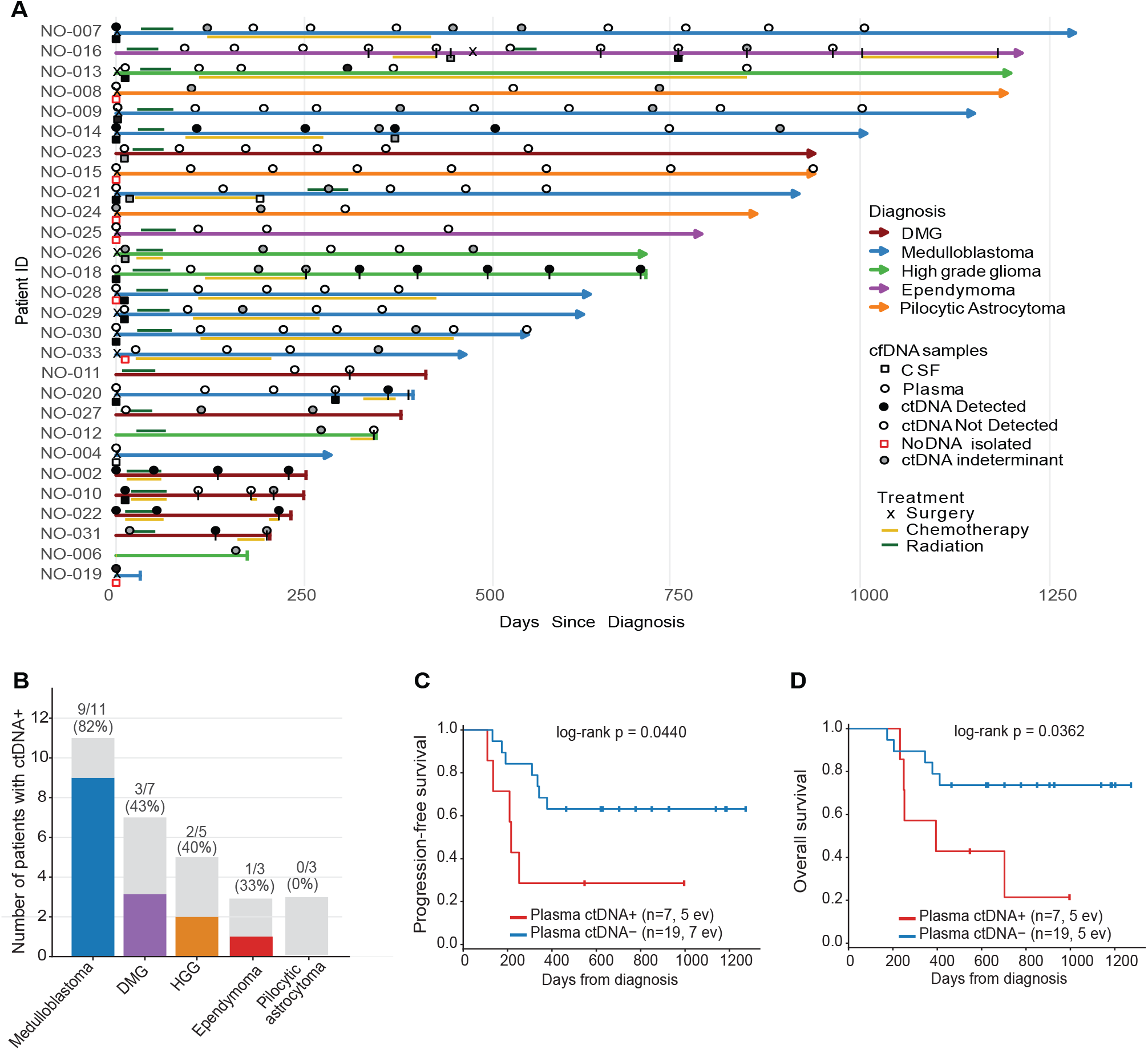
Longitudinal Plasma ctDNA Detection and Clinical Outcomes in Pediatric CNS Patients. **A.** Swimmer plot summarizing treatment timelines and sample collection. Each line represents an individual patient, color-coded by diagnosis. Arrows indicate that patient is alive at last follow-up. A vertical line denotes patient is deceased. Thin dark green line denotes time receiving radiation. Thin orange line denotes time receiving chemotherapy. X symbol on the line indicates surgical resection. Black vertical line on the line indicates disease progression as seen on MRI. Circles and squares mark plasma and CSF samples, respectively, with black filled symbols indicating timepoint of ctDNA positivity. **B.** Proportion of patients with at least one plasma or CSF ctDNA positive sample by diagnosis. **C.** Kaplan–Meier analysis of progression-free survival according to post-diagnosis plasma ctDNA detection. Patients with at least one ctDNA-positive post-diagnosis plasma sample (n = 7) were compared with patients with no ctDNA-positive post-diagnosis plasma samples (n = 19). Survival distributions were compared using the log-rank test (*P* = 0.0440). **D.** Overall survival stratified by post-diagnosis plasma ctDNA positivity.

At the time of diagnosis (Day 0), 6/7 patients with CSF collected (86%) were ctDNA-positive compared to 3/14 (21%) by plasma, though this difference did not reach statistical significance in the paired analysis (McNemar P = 0.062), likely due to the limited number of patients with same-day paired samples (**Supplemental Figure 3A**). The better performance of CSF samples was consistent across subsequent timepoints: CSF positivity rates exceeded plasma at pre-treatment (75% vs. 17%; McNemar P = 0.250), any post-diagnostic timepoint (73% vs. 27%; McNemar P = 0.125), and across all samples from day 0 onward (80% vs. 32%; McNemar P = 0.031). The higher detection rate in CSF reflected consistently greater signal intensity across all assay modalities, with per-patient maximum VAF, ichorCNA tumor fraction, short-to-long fragment ratio, and composite total score each significantly higher in CSF than plasma (all Wilcoxon P ≤ 0.03; **Supplemental Figure 3B**).

Additionally, ctDNA detection rates varied by tumor type (**Figure 4B**). Medulloblastoma had the highest proportion of patients with at least one ctDNA-positive sample (9/11, 82%), followed by DMG (3/7, 43%) and HGG (2/5, 40%). The ependymoma patients had ctDNA detected in 1/3 (33%) patients.

Kaplan–Meier analysis demonstrated that patients with plasma ctDNA detected in any sample collected after the day of diagnosis (day 0; n = 7) had significantly worse progression-free survival (PFS) than patients without detectable plasma ctDNA at any post-diagnosis timepoint (n = 19; log-rank p = 0.044; **Figure 4C**). A similar association was observed for overall survival (OS; log-rank p=0.036; **Figure 4D**). Among patients with post-diagnosis CSF samples available, a trend toward shorter PFS was observed in CSF ctDNA-positive patients, though it did not reach significance (log-rank p=0.168; **Supplementary Figure 4A**). The OS analysis was limited by the absence of events in the CSF ctDNA-negative group (log-rank p=0.370; **Supplementary Figure 4B**).

Cox proportional hazards modeling was used to assess the independent prognostic contribution of plasma ctDNA positivity. Tumor subtype was included as a covariate because DMG and HGG patients have inherently worse survival outcomes and higher plasma ctDNA shedding than other diagnostic groups in this cohort. However, the PFS association remained significant after adjustment (HR 4.52, 95% CI 1.05–19.40, p = 0.027), indicating that ctDNA positivity carries prognostic value independent of subtype. The OS association was attenuated following adjustment (p = 0.109), suggesting the OS signal is at least partly explained by subtype composition (**Supplementary Figure 4C**).

### Longitudinal Plasma ctDNA Mirrors Clinical Progression in a Patient with High-Grade Glioma

To illustrate the utility of longitudinal ctDNA monitoring, a 16-year-old male (NO-018) with an unresectable thalamic high-grade glioma underwent radiation therapy followed by targeted Osimertinib therapy for an EGFR mutation. Despite aggressive therapy, the patient ultimately succumbed to disease approximately two years after diagnosis. CSF and plasma were obtained at the time of diagnosis and several subsequent plasma samples were taken throughout the patient’s clinical course (**Figure 5A**). Radiographically, the patient continued to show signs of progression at each subsequent MRI (**Figure 5B**). CNA of the tumor at diagnosis showed distinct gains in chromosomes 1,2,7 and 9 and near-complete tumor cellularity (ichorCNA tumor fraction 0.9989), although no longitudinal CSF or plasma samples demonstrated matching CNA profiles (**Figure 5C**). Tumor-specific variants, namely *EGFR* p.A244V and *C12orf43* 3’UTR, were absent in early samples obtained during the time of radiation and Osimertinib treatment but found at later plasma timepoints coinciding with radiographic progression later in his clinical course (**Figure 5D**). This emergence of plasma ctDNA was accompanied by a striking rise in total cfDNA yield, which increased from a stable baseline of approximately 5–9 ng to a peak of approximately 22 ng at day 575 before declining slightly at day 696 (**Figure 5E**). The patient died at day 703. Together, these findings demonstrate that rising plasma ctDNA signal, both in variant VAF and total cfDNA yield, can potentially signal tumor progression complementing radiographic evidence.

**Figure 5.**
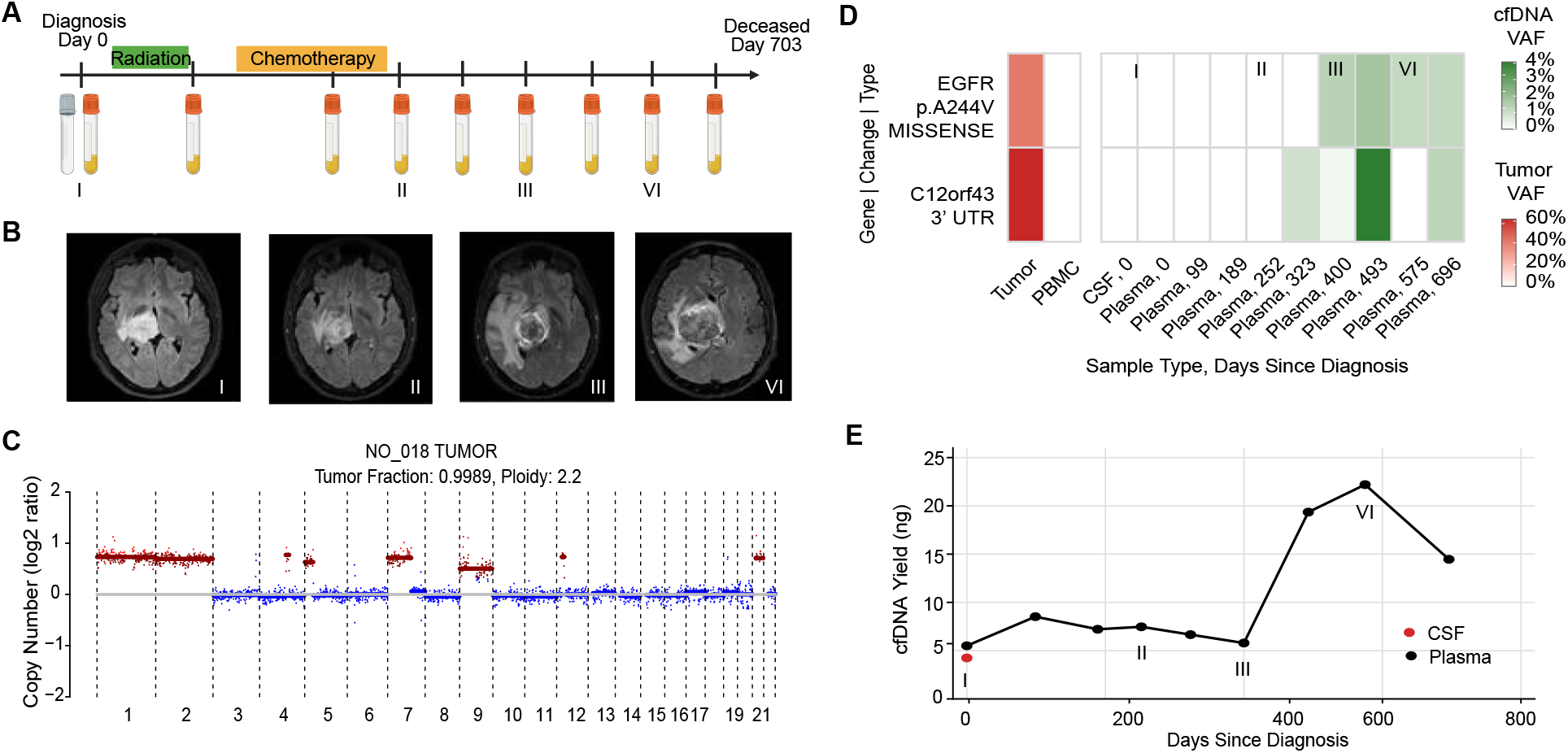
Impact of Clinical Changes on ctDNA Detection. **A.** Clinical time course with radiation and chemotherapy including time points of CSF and blood collection for patient NO_018. **B.** Axial FLAIR MRI series demonstrating progression of the tumor at each subsequent MRI image. **C.** Copy number alteration tumor profile generated by ichorCNA. **D.** Heatmap demonstrating mutational landscape in ctDNA samples for oncogenic tumor-specific variants. Color intensity represents variant allele frequency (VAF), with red indicating VAF in the tumor specimen and green indicating VAF in cfDNA samples. **E**. Line graph depicting the trajectory of extracted cfDNA yield across clinical timeline.

### CSF ctDNA Profiles Tumor at Diagnosis While Rising Plasma Signal Shows Molecular Evidence of Leptomeningeal Dissemination Missed by Imaging

Patient NO-010 was a 17-year-old female diagnosed with diffuse midline glioma (DMG) who received concurrent temozolomide and radiation followed by Avapritinib in clinical trial but died 249 days after diagnosis (**Figure 6A**). The patient had one CSF sample and serial plasma samples drawn throughout her clinical course. Multiple tumor-specific variants ranging from 30 to 92% were found in the CSF sample taken on post-op day 12 via diagnostic lumbar puncture. Those same variants were absent from all the plasma samples (**Figure 6B**). The short-to-long fragment ratio in CSF (1.37) was markedly elevated above the positivity cutoff of 0.381, while plasma samples at days 12, 109, and 179 remained below threshold (0.26, 0.25, and 0.27, respectively; **Figure 6C**).

**Figure 6.**
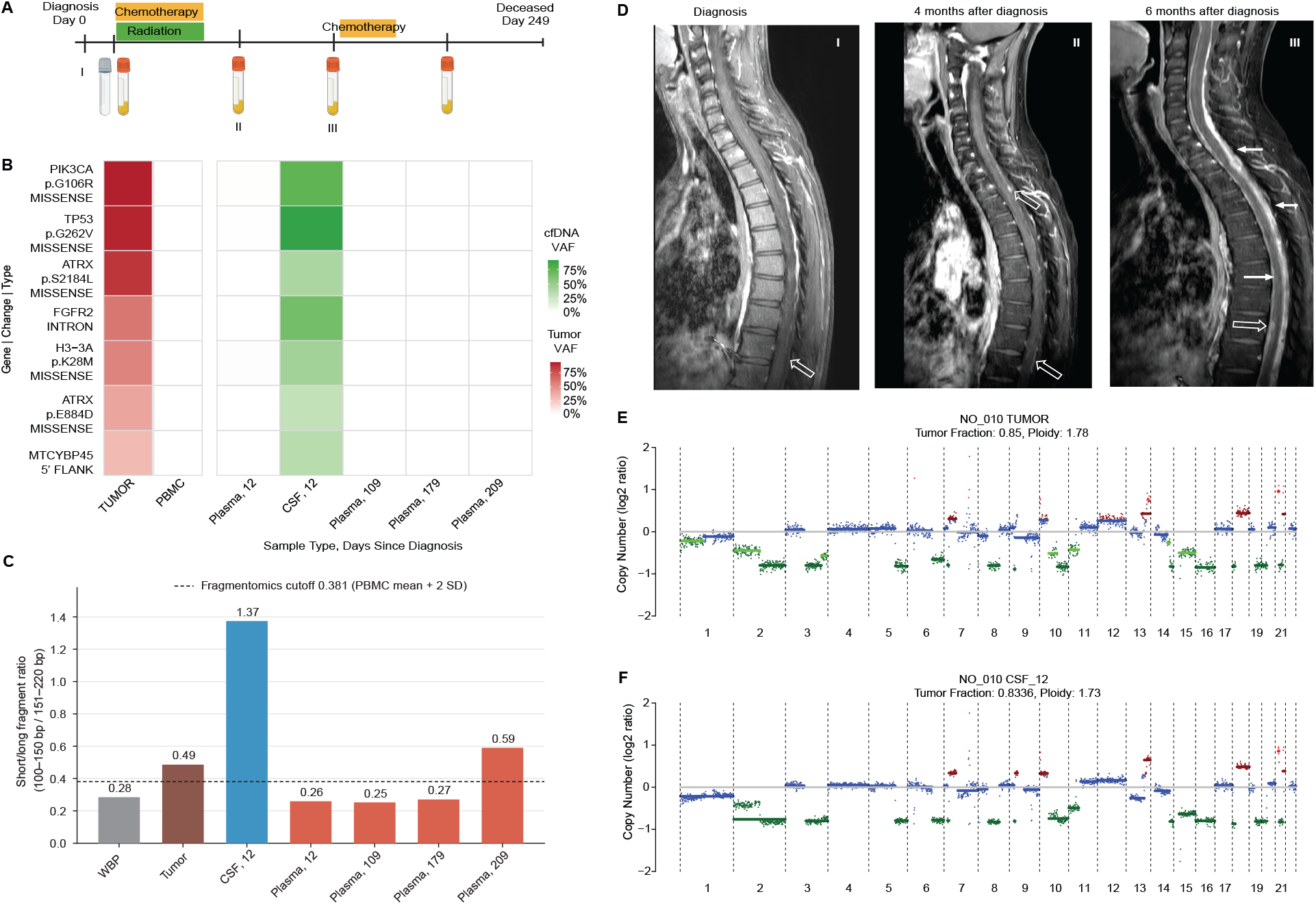
Multimodal ctDNA Characterization: **A.** Clinical time course with radiation and chemotherapy including time points of CSF and blood collection for patient NO_010. **B.** Heatmap of tumor unique variants in longitudinal cfDNA samples. Color intensity represents variant allele frequency (VAF), with red indicating VAF in the tumor specimen and green indicating VAF in cfDNA samples. **C.** Short-to-long fragment (S/L) ratio analysis comparing WBP, tumor, CSF, and plasma samples. Dashed line corresponds to cutoff value which is calculated by taking non-tumor reference (WBP) ratio plus 2 standard deviations. **D.** Sagittal postcontrast T1-weighted MRI image demonstrating no leptomeningeal disease at diagnosis (I). Four months later (II) imaging demonstrating a small nodular and plaque-like enhancing lesions along the dorsal surface of the spinal cord at multiple cervical and thoracic levels (white arrows) consistent with interval disease progression. Six months after diagnosis (III) imaging demonstrates thick plaque-like enhancement coating the ventral and dorsal surfaces of the spinal cord and filling the thecal sac (white arrows). Multiple enhancing intradural-intramedullary nodular lesions are also present (white arrow), consistent with progressive leptomeningeal metastatic disease. **E.** Copy number alteration profiles generated by ichorCNA for tumor sample. **F.** Copy number alteration profiles generated by ichorCNA for tumor positive CSF sample.

Strikingly, the plasma S/L ratio at day 209 rose to 0.59, exceeding the positivity cutoff, coinciding with the development of leptomeningeal metastases visible on spinal MRI at timepoints II and III (**Figure 6D**). Additionally, the day 12 CSF sample demonstrated a tumor fraction of 0.8336, closely mirroring the tumor CNA profile (tumor fraction 0.85, **Figures 6E–F**). Taken together, this case demonstrates that CSF ctDNA can provide comprehensive multimodal tumor characterization before radiographic evidence of dissemination, and that the emergence of plasma ctDNA signal may reflect leptomeningeal dissemination in patients with CNS tumors.

## DISCUSSION

This study demonstrates the feasibility of multimodal, longitudinal ctDNA profiling in a cohort of pediatric patients with CNS tumors using matched plasma and CSF samples. As expected from the anatomical proximity of CSF to the tumor microenvironment, we found stronger ctDNA intensity in CSF samples compared to plasma. Similarly, higher-grade tumors, including DMG, high-grade glioma, and medulloblastoma, demonstrated higher rates of ctDNA positivity than lower-grade tumors such as pilocytic astrocytoma. These observations mirror previous reports linking ctDNA detection to more aggressive tumor biology, higher proliferative activity, and increased genomic instability ^1,12^. Importantly, ctDNA positivity in post-diagnosis plasma samples was significantly associated with shorter progression-free survival and overall survival, consistent with reports that detectable plasma ctDNA identifies patients at elevated risk of progression and poor survival outcomes^8^.

Tumor-informed liquid biopsy approaches, in which somatic variants identified from matched tumor and germline sequencing define a patient-specific target set for cfDNA genotyping, have emerged as a sensitive strategy for ctDNA detection across cancer types^15,22^. By restricting analysis to variants confirmed to be tumor-derived and absent from the germline, this approach substantially reduces the false positive burden arising from clonal hematopoiesis and sequencing noise. A key feature of our variant allele scoring scheme is the differential weighting of variants by their predicted functional consequence. Thus, a score of 1 was used for low-VAF non-coding variants, while a score of 2 required either a pathogenic variant detected at any VAF or a non-coding variant at ≥1%, a threshold widely supported as a boundary between true signal and background noise in plasma ctDNA studies^15,23,24^.

In previous studies, a cut-off value from 3-10% for ichorCNA tumor fraction has been used to determine tumor signal in cfDNA samples^2,20,25^. For our copy number scoring, we used a conservative tumor fraction threshold of >5%, combined with a requirement that it exceed the matched germline estimate by more than 3 percentage points, based on the distribution of tumor fraction observed in germline samples (0-4.1%). For fragmentomic scoring, the short-to-long fragment ratio cutoff was defined empirically as the germline mean plus two standard deviations (0.381), with no matched germline samples exceeding this threshold. We defined short fragments as fragments ranging from 100-150bp based on previous fragmentomic analyses^21,26–28^. Fragmentomics was capped at a maximum score of 1, as elevated short-to-long fragment ratios can reflect non-tumor biology including inflammatory states, infection, or hepatic dysfunction^29,30^.

A central contribution of this work is the proposal of a formalized, multimodal scoring framework for ctDNA detection in pediatric CNS tumors. Rather than relying on any single assay, our composite scoring system integrates orthogonal evidence from targeted sequencing, copy number profiling, and fragmentomics into a reproducible positivity call that, to our knowledge, has not been previously used for pediatric CNS tumors. The differential weighting of variants by functional consequence, the germline-anchored thresholds for CNA and fragmentomic scoring, and the requirement for a composite score of ≥2 to define positivity each represent deliberate choices that reduced false positive calls while maintaining sensitivity across a heterogeneous cohort. The integration of this composite scoring methodology demonstrably improved ctDNA detection beyond what any single modality achieved. Targeted sequencing alone identified 26 ctDNA-positive samples, while CNA alone identified 20; combining all three modalities increased this to 38 positive samples, representing a meaningful gain in sensitivity. For example, two medulloblastoma patients in our cohort lacked any tumor-specific mutations hindering our ability to confidently track the mutational landscape in the patient’s cfDNA. In those patients, the use of copy number analysis allowed us to find positive ctDNA signal in subsequent longitudinal samples. While fragmentomic analysis did not identify additional positive samples independently, it did frequently confirm ctDNA positivity. The greatest benefit was seen in CSF samples, where fragment size profiles often supported the presence of ctDNA detected by targeted sequencing or copy number profiling. These findings suggest that fragmentomics may be used to improve confidence in ctDNA classification.

At the same time, the thresholds determining this scoring framework were defined internally and validated in a single cohort, which awaits generalizability until tested in independent cohorts. The 1% VAF boundary, the 5% tumor fraction cutoff, and the germline mean + 2 SD fragmentomic threshold were each grounded in the distribution of our matched germline samples, but may require recalibration across sequencing platforms, input quantities, and nuances in patient populations. More broadly, tumor-informed multimodal ctDNA analysis requires matched tumor tissue, germline, and longitudinal cfDNA samples for each patient which can prove to be a logistical challenge limiting scalability. The requirement for all three sample types means that loss of any single component can render a patient unevaluable.

Future work should prioritize prospective validation of composite scoring frameworks in larger, multi-institutional cohorts. Establishing such harmonized criteria is a prerequisite for translating multimodal liquid biopsy from a research tool into a clinically actionable assay. Beyond analytic considerations, opportunities for serial CSF collection were limited by clinical availability, reducing statistical power. Extracting cfDNA from a subset of CSF samples proved technically challenging due to low input quantities. The resulting diminished sequencing coverage in these samples restricted our ability to draw definitive conclusions regarding the relationship between ctDNA detection and the method of CSF collection. Finally, although CSF offered clearly superior analytical sensitivity, serial plasma collection proved far easier to integrate into routine visits, reinforcing the need to improve plasma-based detection strategies for longitudinal disease monitoring.

Future studies with this dataset will involve integrating ctDNA detection rates with radiographic tumor volumetrics, treatment exposures, and clinical outcomes. Machine learning approaches utilizing DNA sequencing with a classification-based algorithm are underway to add an additional technique to the multimodal methodology. Our findings demonstrate that longitudinal plasma and CSF collection is feasible within routine clinical workflows and support the continued development of multimodal ctDNA assays as a tool for diagnosis, disease monitoring, and early detection of progression in pediatric CNS tumors.

## ACKNOWLEDGEMENTS

This research includes work performed in the TGen Collaborative Sequencing Center, a City of Hope Comprehensive Cancer Center supported shared resource (NCI-P30CA033572). Large language models were used in the preparation of this manuscript to improve grammatical accuracy and readability. The authors reviewed all edits and retain sole responsibility for the accuracy and integrity of the content.

## AUTHORSHIP STATEMENT

SG, SM, FPB, and RM contributed to study conception and design. SG, BP, and SM coordinated sample collection and inventory across clinical and research sites. AM, TCC, AH, SK, and JK conducted sample processing and sequencing. SM conducted all bioinformatics analyses. SG and SM performed data interpretation, figure generation, and manuscript writing. FB and RM supervised the study. All authors reviewed and approved the final manuscript.

## DATA AVAILABILITY

Processed variant calls can be found on Zenodo under 10.5281/zenodo.23743662. Segmentation files from ichorCNA have been divided into two Zenodo repositories due to storage constraints and can be found in 10.5281/zenodo.23896717 (patients 002-021 cfDNA) and 10.5281/zenodo.23896810 (patients 021-051 cfDNA and tumor/normals). Raw sequencing data may be made available from the corresponding author upon reasonable request and with appropriate institutional approvals.

## SUPPLEMENTARY FIGURES

**Supplementary Figure 1:**
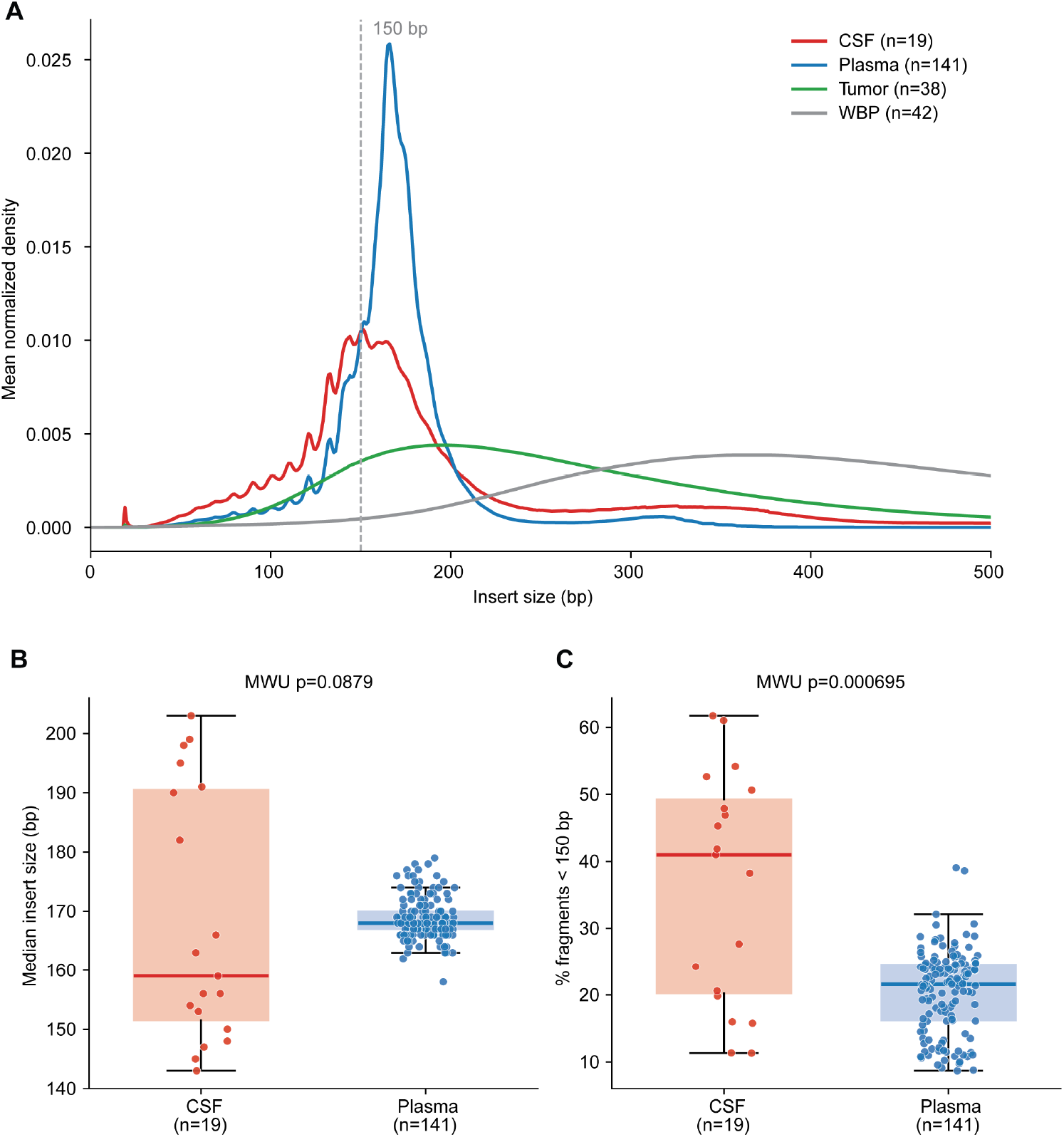
Fragmentomic analysis across 28 patients. **A.** Density plot showing distribution of fragments from 0-500bp. **B.** Shorter median insert size for CSF samples compared to plasma. **C.** Significantly more short fragments (<150bp) present in CSF samples compared to plasma.

**Supplementary Figure 2:**
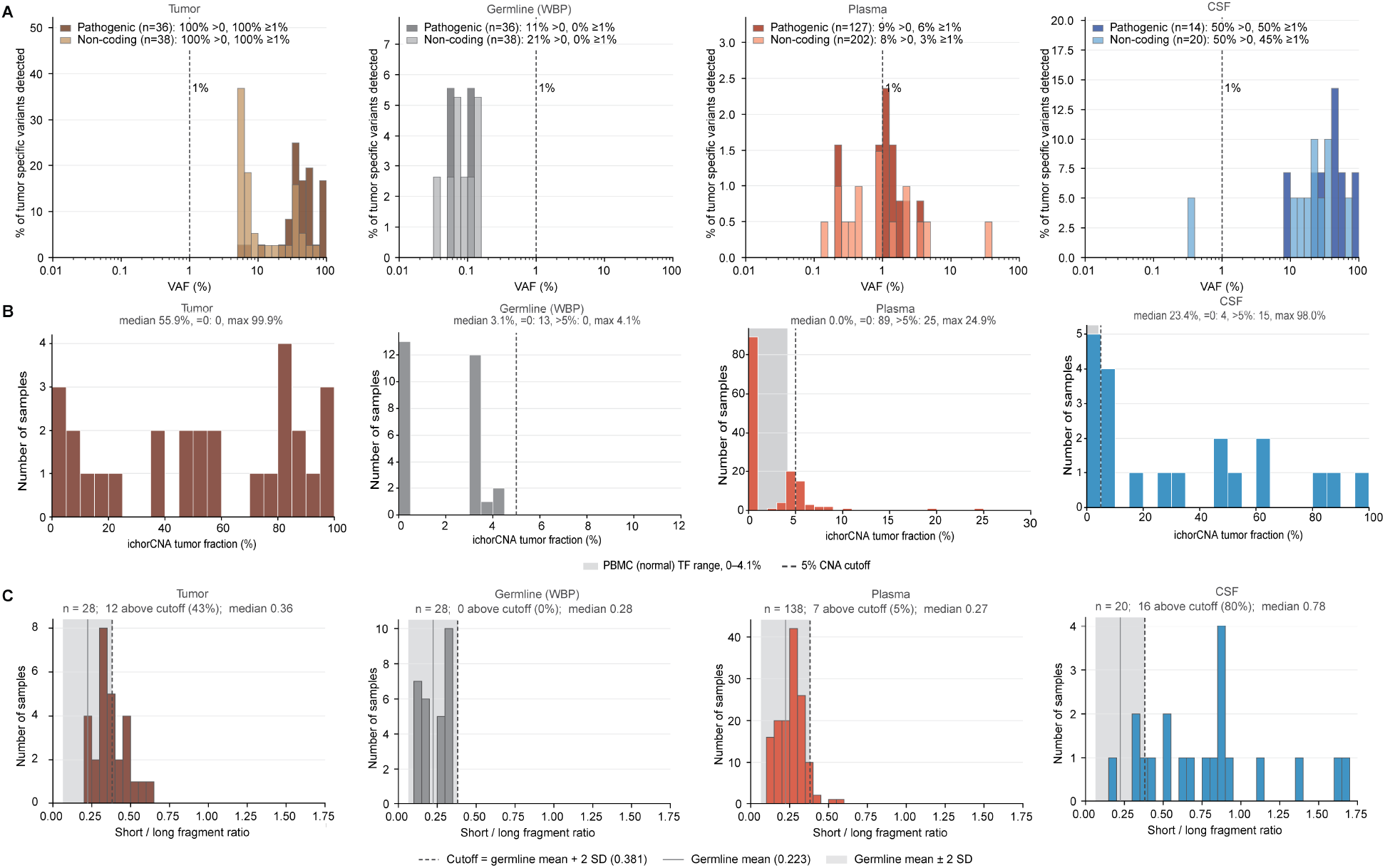
Derivation of tumor positivity using WBP samples. **A.** Distribution of tumor specific VAF for tumor, germline, plasma and CSF samples. Variant allele frequency (ALT reads / total depth) in tumor, matched germline, plasma and CSF, at positions of tumor-confirmed somatic variants, shown separately for pathogenic driver variants (dark) and non-coding variants (light; intronic, UTR, flanking or silent). **B.** Distribution of ichorCNA tumor fraction for tumor, germline, plasma and CSF samples. VAF distribution at tumor-specific variant positions by sample type. **C.** Short/long fragment ratio distribution for tumor, germline, plasma and CSF samples.

**Supplementary Figure 3:**
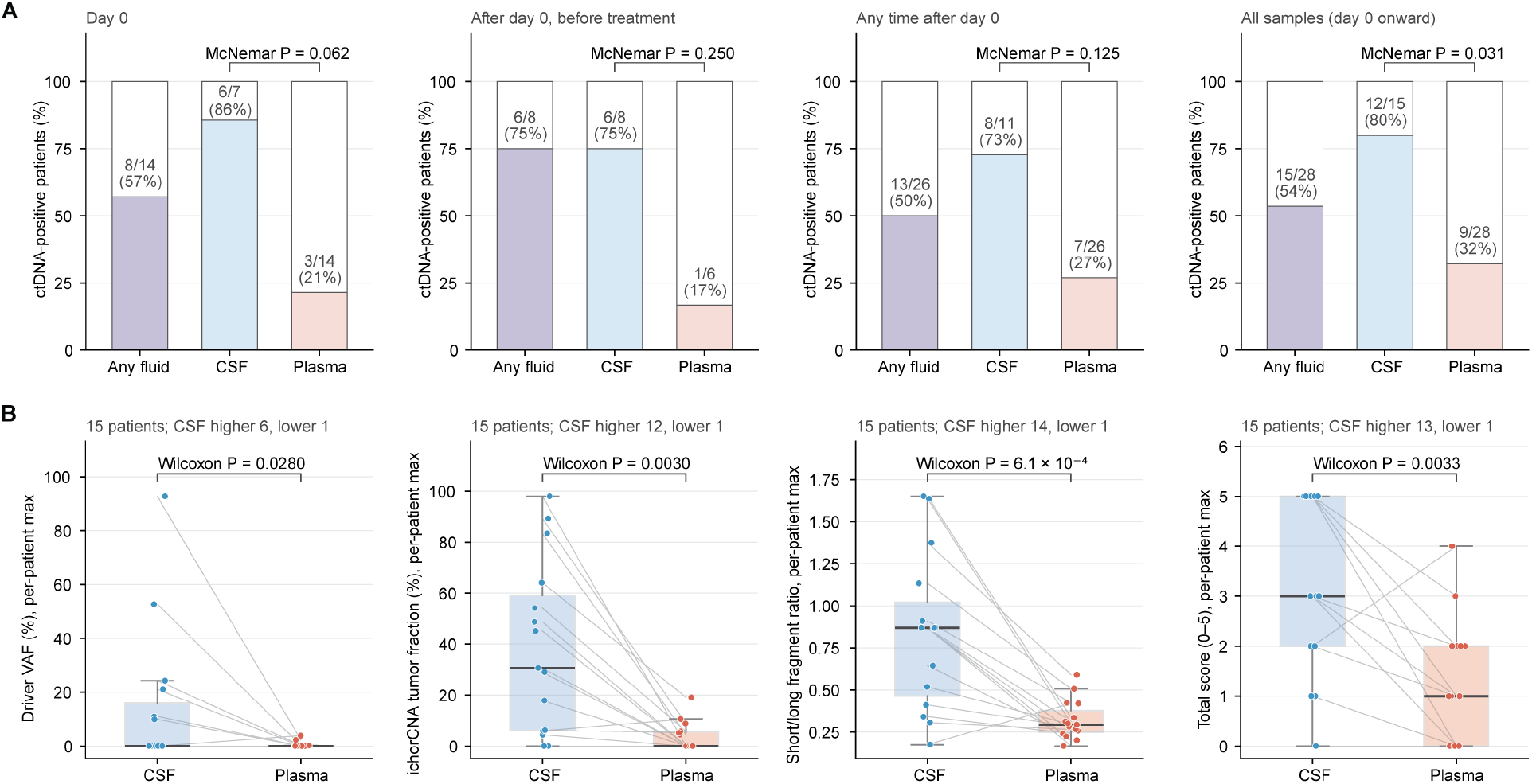
Within Patient Biofluid Comparison. **A.** Percentages of ctDNA positive patients in any fluid, CSF or plasma at different timepoints. **B.** Comparisons of different tumor metrics and composite scoring within the two biofluids.

**Supplementary Figure 4:**
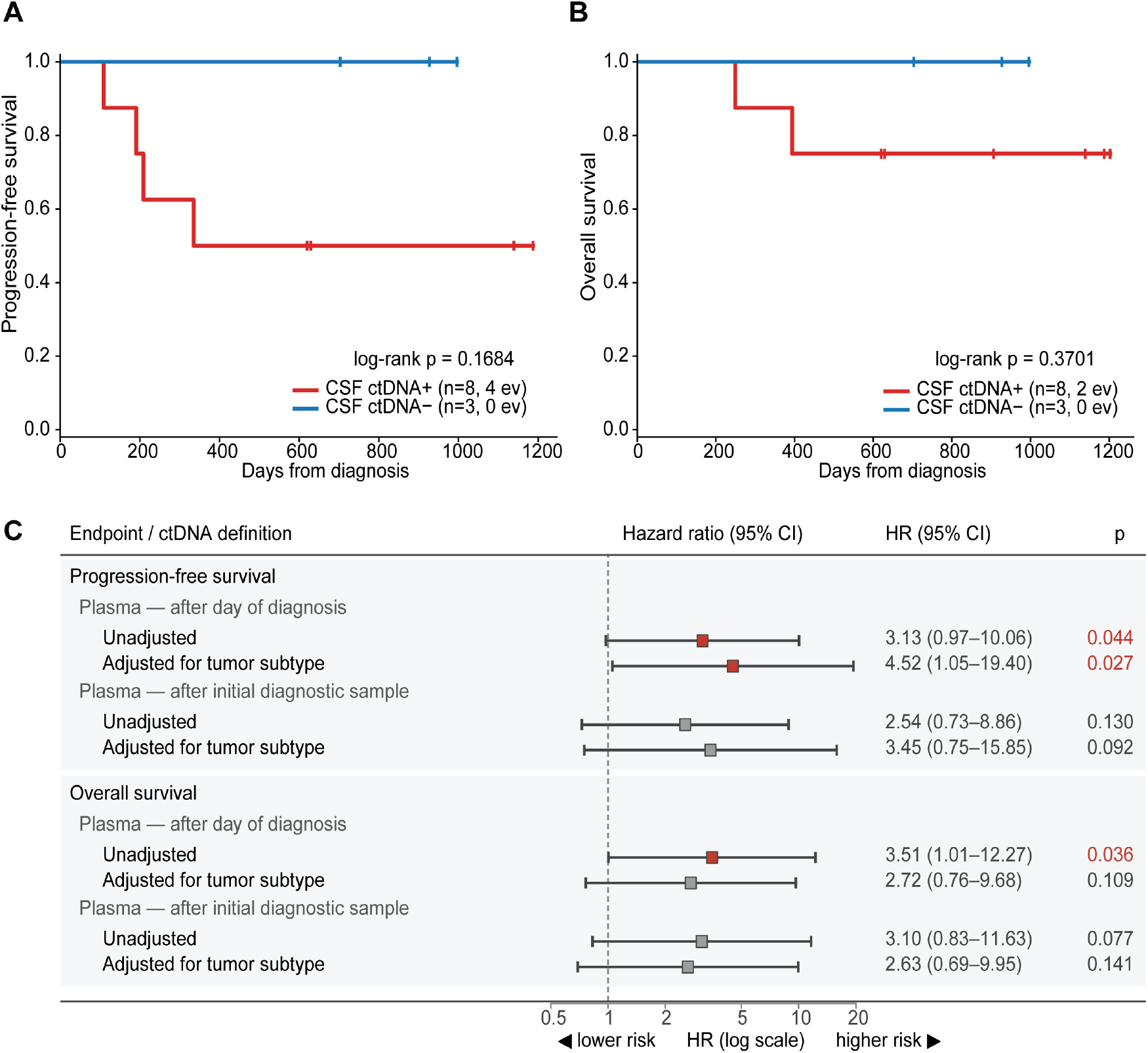
Post-Diagnosis ctDNA Status and Survival by Biofluid Source. **A.** Kaplan–Meier curves for progression-free survival in patients with at least one post-diagnosis CSF sample (n=11; 8 ctDNA+, 4 progression events; 3 ctDNA−, 0 events). Log-rank p=0.168. **B.** Overall survival in the same patients (n=11; 8 ctDNA+, 2 deaths; 3 ctDNA−, 0 deaths). Log-rank p=0.370. Hazard ratio for overall survival was not estimable because no CSF ctDNA-negative patient experienced an event. **C.** Forest plot of unadjusted and tumor subtype-adjusted hazard ratios (95% CI) for progression-free survival (n=26, 12 events) and overall survival (n=26, 10 events), comparing patients with at least one post-diagnosis plasma ctDNA-positive sample (n=7) versus those without (n=19). P values are from log-rank tests; adjusted models were stratified by tumor subtype.

**Supplementary Figure 5:**
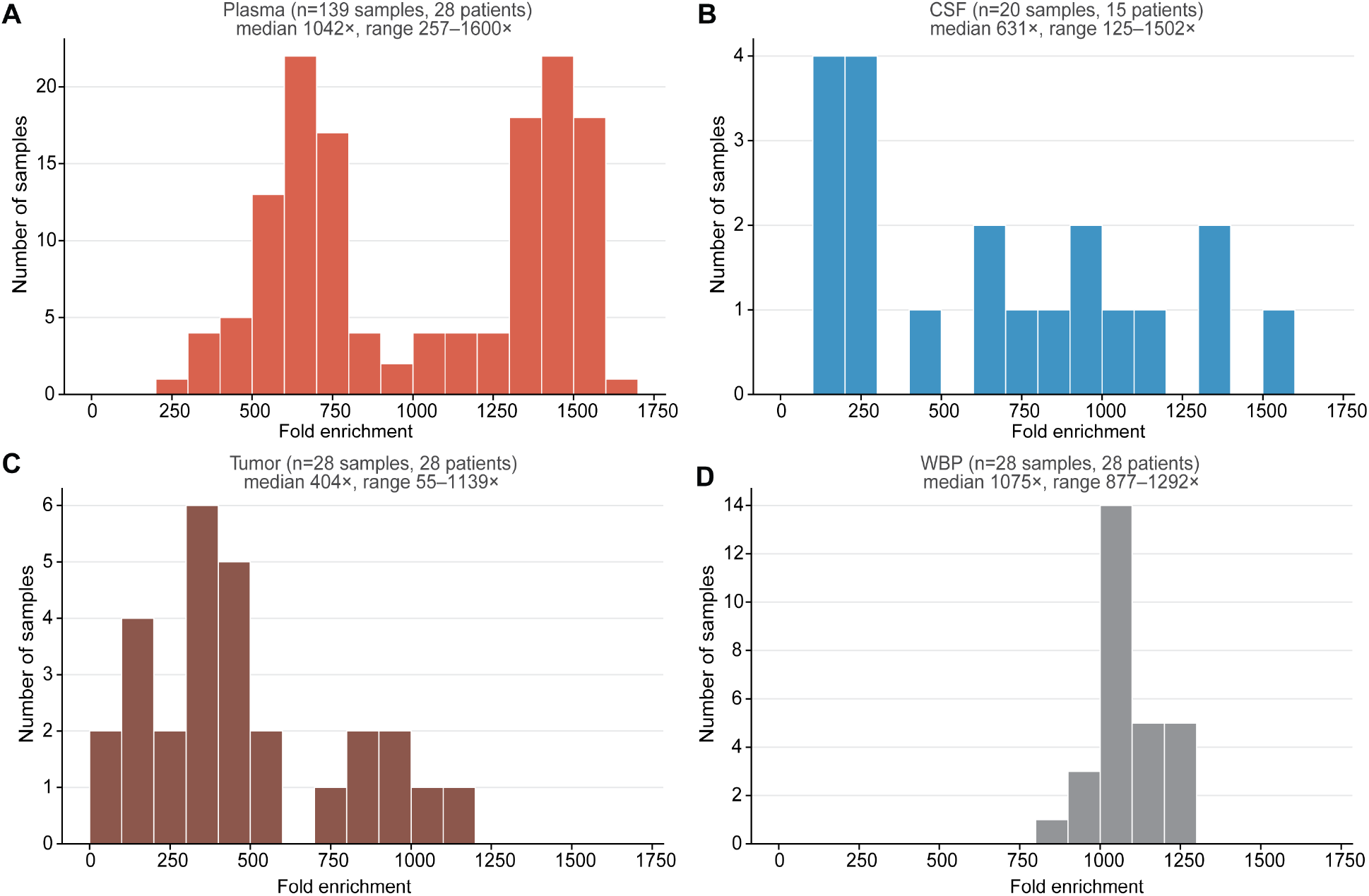
Targeted Sequencing Coverage and Fold Enrichment Metrics. **A.** Plasma, **B.** CSF, **C.** Tumor and **D.** Whole Blood Pellets.

